# The dopamine receptor D_5_ gene shows signs of independent erosion in Toothed and Baleen whales

**DOI:** 10.1101/556035

**Authors:** Luís Q. Alves, Juliana Alves, Rodrigo Ribeiro, Raquel Ruivo, L. Filipe C. Castro

## Abstract

To compare gene *loci* considering a phylogenetic framework is a promising approach to uncover the genetic basis of human diseases. Imbalance of dopaminergic systems is suspected to underlie some emerging neurological disorders. The physiological functions of dopamine are transduced via G-protein-coupled receptors, including DRD_5_ which displays a relatively higher affinity towards dopamine. Importantly, DRD_5_ knockout mice are hypertense, a condition emerging from an increase in sympathetic tone. We investigated the evolution of DRD_5_, a high affinity receptor for dopamine, in mammals. Surprisingly, among 124 investigated mammalian genomes, we found that Cetacea lineages (Mysticeti and Odontoceti) have independently lost this gene, as well as the burrowing *Chrysochloris asiatica (Cape golden mole).* We suggest that DRD_5_ inactivation parallels hypoxia-induced adaptations, such as peripheral vasoconstriction required for deep-diving in Cetacea, in accordance with the convergent evolution of vasoconstrictor genes in hypoxia-exposed animals. Our findings indicate that Cetacea are natural knockouts for DRD_5_ and might offer valuable insights into the mechanisms of some forms of vasoconstriction responses and hypertension in humans.

## 1. Introduction

Dopamine is a neurotransmitter essential for brain function, regulating various physiological processes including locomotion, cognition, and neuroendocrine functions [1,2]. Dopamine molecular actions are transduced *via* a specific group of G-protein coupled receptors entailing two major classes: DRD_1_-like and DRD_2_-like receptors [3,4]. While DRD_1_-like receptors stimulate cAMP production postsynaptically, DRD_2_-like receptors inhibit cAMP production both pre and postsynaptically [3]. The genomic structure of the underlying genes is also distinct, with DRD_1_-like receptors yielding single exon coding regions [3]. DRD_1-2_ receptor classes have diversified in vertebrate evolution most likely as a result of genome duplications [4]. Interestingly, agonist and antagonist amino acid site conservation suggests evolutionary stasis of dopaminergic pathways [4]. Among DRD_1_-like receptors, the DRD_5_ subtype displays distinctive features, namely a relatively higher affinity towards dopamine, a putative agonist-independent activity and low level, yet widespread, brain expression [3,5–7]. Nonetheless, the DRD_5_ seems to display distinct regional and cellular distribution patterns in the brain, when compared to the DRD_1_ and DRD_2_ subtypes, with protein enrichment detected in the cerebral cortex, hippocampus and basal ganglia [7,8]. Peripheral expression has also been found in the adrenals [9], kidney [10] and gastrointestinal tract [11]. Despite the association with schizophrenia [12], attention-deficit/hyperactive disorder [13] and substance abuse [14], gene targeting studies revealed that DRD_5_ knock-out mice develop hypertension, showing increased blood pressure from 3 months of age [2]. This hypertensive phenotype appears to result from a central nervous system defect, leading to an increase in sympathetic tone and, consequently, vasoconstriction [2]. Besides neuronal impairment, DRD_5_ disruption was also suggested to counter-regulate and modulate the expression of the prohypertensive Angiotensin II Type 1 Receptor (AT_1_R), involved in renal salt balance, blood pressure and vasoconstriction [15–17].

Whole genome sequencing has greatly expanded our capacity to comprehend evolutionary history, the role of adaptation or the basis for phenotype differences across the tree of life. Multigenome comparisons have also been powerful to recognize the molecular basis of human diseases, a field named as phylomedicine [18–21]. Here we investigate the evolution of DRD_5_ in mammalian species. By analysis 124 genomes covering 16 orders, we show that independent coding debilitating mutations occurred in the ancestors of Mysticeti and Odontoceti (Cetacea). Our findings suggest that these species are natural KOs for this dopamine receptor and might offer valuable insights into the mechanisms of some forms of essential hypertension.

## 2. Results

To examine the annotation tags and distribution of the DRD_5_ gene across mammals, 119 annotated mammalian genomes available at NCBI (National Center of Biotechnology Information), were scrutinized for the presence of DRD_5_ gene annotation and each respective protein product description screened for the ‘low-quality protein’ (LQ) tag. This examination resulted in 10 species presenting DRD_5_ annotation tagged as ‘low-quality protein’ producing gene, including *Ovis aries* (sheep), *Phascolarctos cinereus* (koala), *Bison bison bison* (plains bison), *Myotis davidii* (vesper bat), *Ochotona princeps* (American pika) and 5 cetacean species. The latter included *Lagenorhynchus obliquidens* (Pacific white-sided dolphin), *Neophocaena asiaeorientalis asiaeorientalis* (Yangtze finless porpoise), *Delphinapterus leucas (beluga whale), Physeter catodon (sperm whale) and Orcinus orca* (killer whale). Each genomic sequence corresponding to the DRD_5_ LQ annotations was examined and the coding sequence (CDS) manually predicted [22]. Given the prominence of DRD_5_ LQ annotations in Cetacea we scrutinized other cetacean species with available, but unannotated genomes, *Balaenoptera bonaerensis* (Antarctic minke whale), *Eschrichtius robustus* (gray whale), *Balaena mysticetus* (bowhead whale), *Sousa chinensis* (Indo-Pacific humpback dolphin), or with annotated genomes lacking DRD_5_ annotations: *Lipotes vexillifer* (Yangtze River dolphin) and *Balaenoptera acutorostrata scammoni* (minke whale) (Figure 1). Additionally, *Tursiops truncatus* (bottlenose dolphin), presenting a seemingly intact DRD_5_ gene annotation, without the ‘low-quality protein’ tag, as well as *Hippopotamus amphibius* (common hippopotamus) predicted DRD_5_ coding sequence (Figure 1), representing the closest extant lineage of Cetacea, were equally inspected. Other mammals with annotated genome without DRD_5_ gene annotation were also scrutinised, namely: *Microcebus murinus* (gray mouse lemur), *Jaculus jaculus* (lesser Egyptian jerboa), *Chrysochloris asiatica* (Cape golden mole), *Erinaceus europaeus* (western European hedgehog), *Elephantulus edwardii* (Cape elephant shrew) and *Condylura cristata* (star-nosed mole). In total, 124 mammalian species were inspected and an in-depth description regarding analysed species list and genomic sequences accession numbers are available at Supplementary Table 1. The Supplementary Figure 1 presents a multiple translation alignment of the NCBI non-’low-quality protein’ tagged mammalian DRD_5_ orthologous sequences. The alignment also includes the predicted *H. amphibius* DRD_5_ sequence and excludes T. truncatus DRD_5_ sequence, afterwards demonstrated as pseudogenized (see below). The examined sequences exhibit a substantial degree of conservation (average pairwise identity of over 80%), with minor variations in expected protein size. The protein sequence conservation is particularly noticeable at the c-terminus (see below).

**Figure 1.**
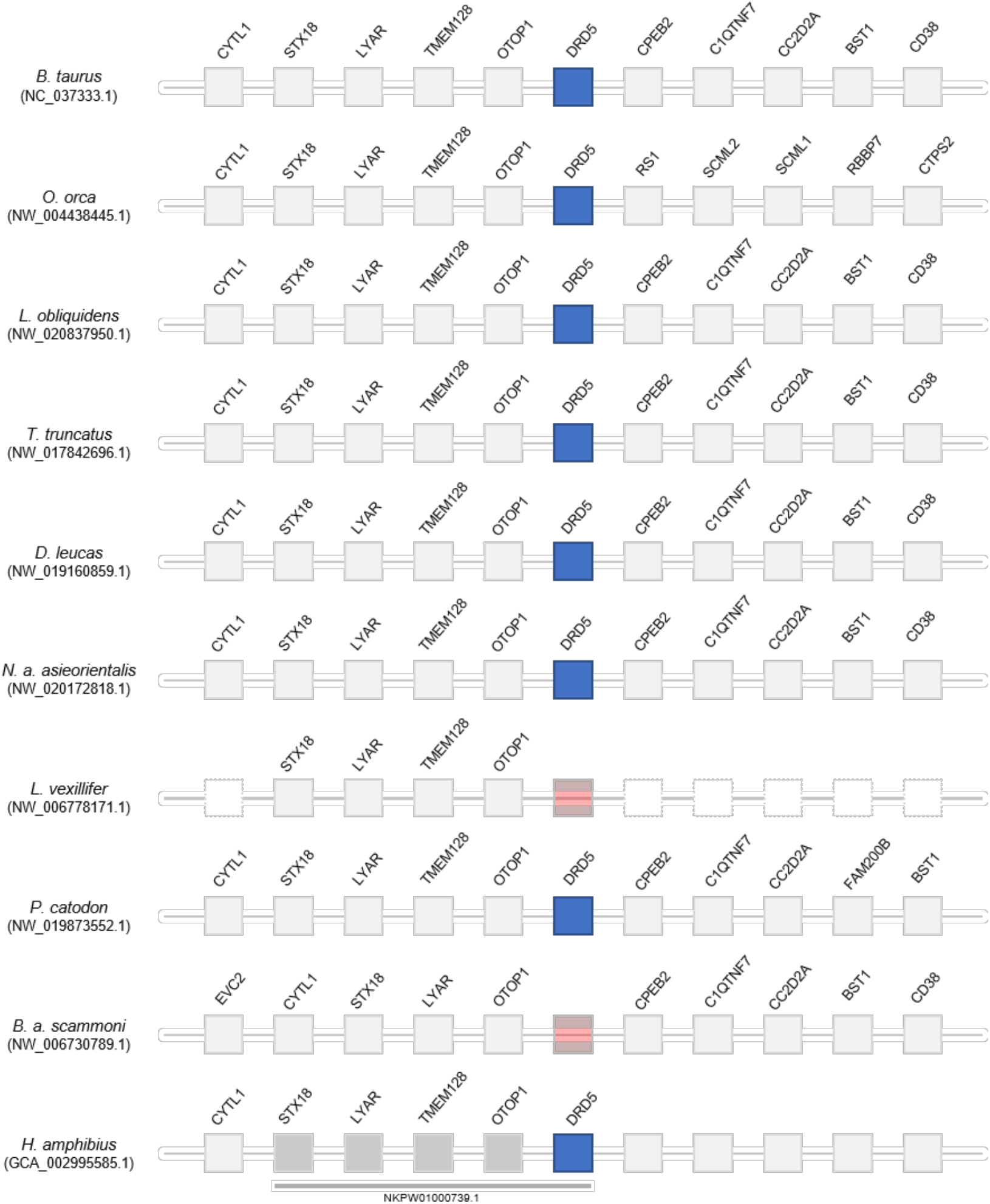
Comparative synteny maps of DRD_5_ genomic *locus* in Cetacea and *H. amphibius.* Blue squares indicate the presence of DRD_5_ in genome annotation. Red squares represent the absence of DRD_5_ annotation in the corresponding genome. *H. amphibius* genes found in the same genomic scaffold are represented by dark grey squares and underlined with a bar indicating the corresponding accession number. Not found genes due to the end of genomic region are represented by white squares.

### 2.1. ORF disrupting mutations of DRD5 in Cetacea

For cetacean species with annotated genomes, we started by examining the DRD_5_ gene locus, including neighbouring genes, to verify and elucidate the orthology of the annotated and nonannotated genes and outline the genomic regions to be inspected. All analysed loci were found to be conserved, including in both L. vexillifer and B. a. scammoni, which lacked previous DRD_5_ gene annotations (Figure 1). Subsequent manual annotation of all collected cetacean genomic sequences revealed DRD_5_ gene erosion across all analysed species, except in B. a. scammoni, for which the DRD_5_ coding status could not be accessed due to fragmentation of the 5’ end of the respective genomic region (presence of sequencing gaps (Ns)). In detail, a conserved 2 nucleotide deletion was detected for the full set of Odontoceti species examined, except for *P. catodon* that presented a premature stop codon in the middle of the gene and a single nucleotide insertion near the end of the gene (Figure 2). The deletion alters the reading frame, leading to a drastic change in downstream amino acid composition. Additionally, a premature stop codon shortens the protein’s expected size—31 amino acid shorter than hippopotamus—a characteristic not found in any of the examined coding DRD5 sequences. Additionally, non-conserved mutations were found in *L. vexillifer* which presented 2 premature stop codons and *D. leiicas* which displayed a single nucleotide deletion near the 5’ end of the gene (Figure 2). *D. leiicas* presumed DRD5 sequence presented another noticeable feature. A massive and abrupt alignment identity decrease was observed when aligned with the *Bos* taurus (cow) reference against the genomic target region of *D. leiicas.* The alignment identity drop was observed approximately in the middle of the complete alignment length for this species, suggesting that the DRD5 gene sequence is interrupted, and further supporting pseudogenization (Figure 3). Regarding Odontoceti, at least one ORF-abolishing mutation was validated using available genomic Sequence Read Archives (SRAs) experiments for all studied species, excluding *S. chinensis* and *L. vexillifer* for which no genomic sequencing runs were available at the NCBI SRA database (Supplementary Material 1).

**Figure 2.**
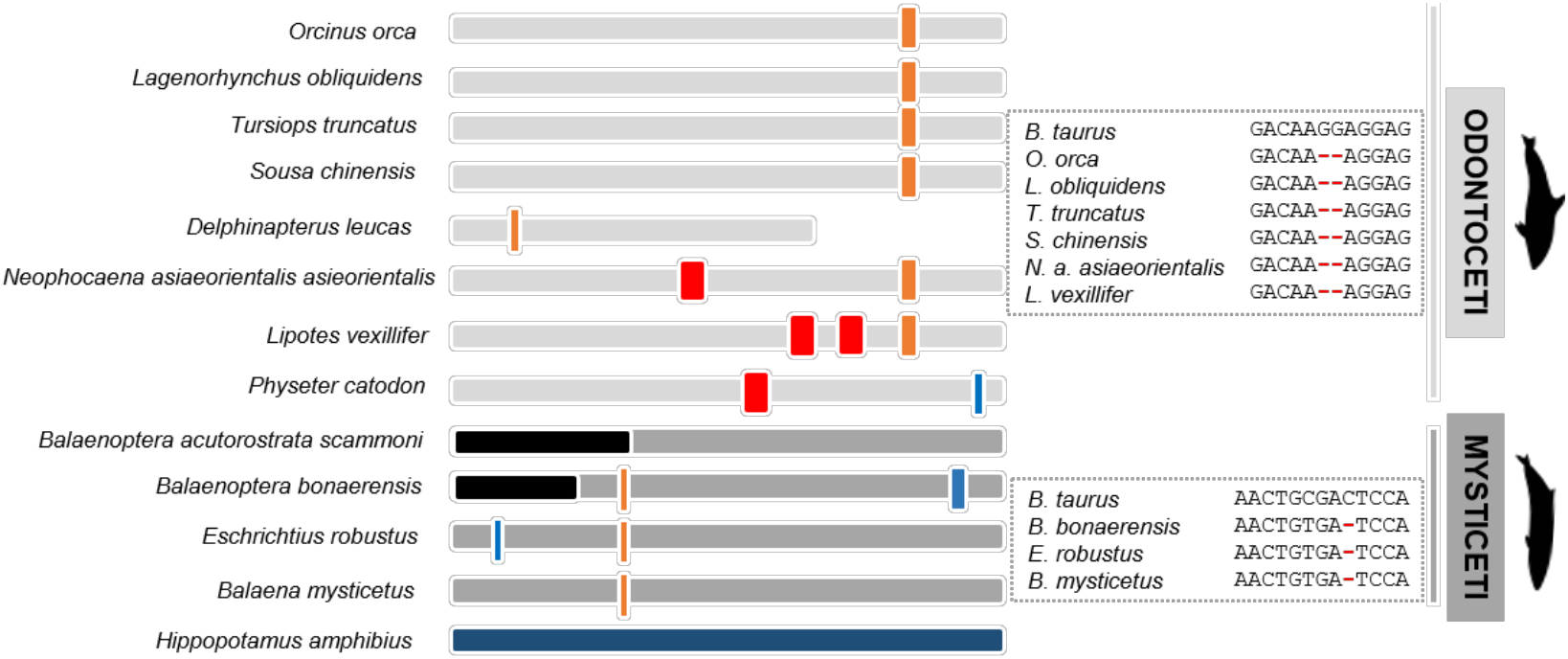
Schematic representation of the DRD5 gene ORF abolishing annotated mutations regarding Cetacea suborders Odontoceti (light gray) and Mysticeti (dark gray). Thicker bars represent 2 nucleotide insertions (blue) or deletions (orange). Thinner bars represent single nucleotide insertion (blue) or deletion (orange). Premature stop codons are represented by red thick bars. Unknown regions, either resulting from genome poor assembly or coverage (Ns in *Balaenoptera acutorostrata scammoni)* or due to small genomic scaffold size (*Balaenoptera bonaerensis)*, are represented by black regions. *Hippopotamus amphibius* presents a complete functional DDR_5_ gene. Two elucidative boxes represent the conserved mutations found in Odontoceti (2 nucleotide deletion) and in Mysticeti (1 nucleotide deletion).

**Figure 3.**
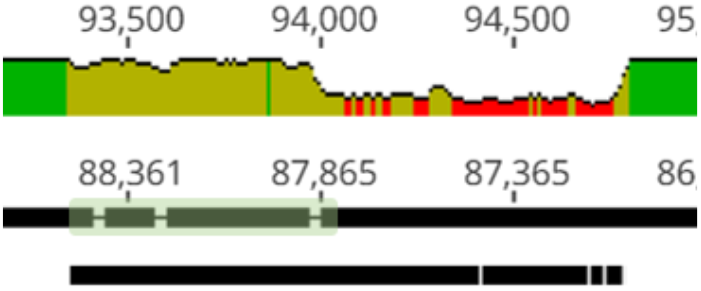
*B. taurus* reference DRD_5_ gene (single-exon) aligned against the genomic target regions of *D. leucas*. Lower bar represents the reference exon and the upper bar represents the genomic target region. An alignment identity graph is presented above the alignment. Green represents very high alignment identity values, followed by yellow (mid to high alignment identity values) and finally red, indicating a very low alignment identity value. The hypothetical approximate DRD_5_ gene real size in *D. leucas* is marked by the clear abrupt alignment identity drop and is represented by a green shadowed rectangle.

Regarding the Mysticeti suborder, a conserved single nucleotide deletion was detected in all species except *B. a. scammoni* (Figure 2). A non-conserved 2 nucleotide insertion was found also found in *B.* bonaerensis and an insertion of one nucleotide was detected at *E. robustus* DRD_5_ sequence (Figure 2). Again, at least one ORF-abolishing mutation was validated using genomic SRA experiments for all analysed species (Supplementary Material 1). Importantly, no conserved mutations were detected between Odontoceti and Mysticeti clades, suggesting that DRD_5_ pseudogenization events occurred independently after the divergence of both lineages (Figure 2). To increase the robustness of our analysis, we further scrutinized the genome of the extant sister clade of the Cetacea, the Hippopotamidae and were able to predict a fully functional CDS for DRD_5_ in *H. amphibius*, supporting the loss of *DRD_5_* after Cetacea diversification.

### 2.2. Other mammalian species showing DRD5 LQ tags have a coding gene

Our initial analysis revealed the presence of at least one ORF-abolishing mutation in *O. aries, P. cinereus*, *B. bison bison* and *O. princeps.* These are in some cases suggestive of gene inactivation and not sequencing artefacts [18,22]. To investigate whether these DRD_5_ gene sequences are eroded in these species, we performed a meticulous manual annotation including SRA validation. Our results revealed sequencing reads supporting the functionality of the gene, rebutting each inactivation mutation and suggesting that DRD_5_ is, in fact, coding in these species (Supplementary Material 2). Regarding *M. davidii*, the fragmentation of the genomic region (Ns) flanked by upstream and downstream DRD_5_ neighbouring genes impeded us to infer the DRD_5_ coding status in this species, although, the visible nucleotide aligning regions showed no ORF disruptive mutations, excepting for an 8 nucleotide deletion near the end of the gene, leading to a STOP codon, suggesting that this gene possibly is lightly truncated but still functional in this species. Interestingly, regarding *O.* princeps mutational SRA validation, two scenarios were observed: approximately 50% of aligned reads supported the absence of a premature stop codon in the DRD_5_ gene of this species, with the remaining set of aligning reads supporting the presence of a premature stop codon in the same species. This suggest that an allelic pseudogenization event might have occurred in *O. princeps.*

### 2.3. Chrysochloris asiatica presents a non-functional DRD5 gene

Next, we examined other mammalian species with annotated genome yet lacking DRD_5_ gene annotations. Results were inconclusive regarding the coding status of *M. murinus, C. cristata, J. jaculus* and *E. europaeus* DRD_5_, due to the fragmentation (Ns) of the genomic region flanked by the upstream and downstream DRD_5_ neighbouring genes, and the unavailability of whole genome shotgun contigs, spanning our target gene. For *E. edwardii* we were able to deduce a fully functional DRD_5_ coding sequence. Curiously, *C. asiatica* presented a single nucleotide insertion in the 5’ end of the gene (validated by genomic SRA, see Supplementary Material 3), suggesting that DRD_5_ is pseudogenized in this species.

## 3. Discussion

The rise of large-scale genomic sequencing projects has emphasized the role of gene loss as a potent driver of evolutionary change: underlying phenotypic adaptations or neutral regressions in response to specific environmental cues and niches [23–28]. Interestingly, gene loss mechanisms seem pervasive in lineages that endured drastic environmental adaptations in the course of evolution, such as Cetacea, entailing niche-specific physiological and morphological adaptations [25,29–33].

By comparing 124 mammalian genomes, we document the independent erosion of a dopamine receptor, DRD_5_, in both Cetacea lineages, Mysticeti and Odontoceti. Dopamine, a neurotransmitter and signalling molecule, is involved in distinct functions both in the central nervous system and peripheral tissues: including movement, feeding, sleep, reward, learning and memory as well as in the regulation of olfaction, hormone pathways, renal functions, immunity, sympathetic regulation and cardiovascular functions, respectively [3]. More specifically, the disruption of DRD_5_-dependent pathways in rodents was shown to increase blood pressure and sympathetic tone, promoting vasoconstriction, thus yielding a hypertensive phenotype [2,15].

The observed gene loss distribution, and predicted phenotypic outcome, is consistent with the peripheral vasoconstriction mechanism described in Cetacea, suggested to counterbalance deep-diving induced hypoxia [34,35]. In fact, Cetacea have developed a number of physiological adaptations to offset hypoxia: notably, increased blood volume and oxygen-transport protein levels (hemoglobin, neuroglobin and myoglobin), allowing oxygen stores in blood, muscle tissues, and brain, reduced heart rate or bradycardia, apnea and peripheral vasoconstriction [34–36]. Peripheral vasoconstriction allows the regional compartmentalization of blood supplies, reducing blood flow in more hypoxia-tolerant tissues, such as skin, muscle, spleen or kidney, while maintaining arterial blood flow to the central nervous system and heart [36]. Moreover, the deviation of muscle blood supply reduces lactate accumulation [36]. Thus, in Cetacea, DRD_5_ loss could contribute for the peripheral vasoconstriction requirements of diving. Also, by shifting renal salt balance, DRD_5_ could also play a role in the maintenance of an adequate blood volume and pressure [15–17,37].

The predicted vasoconstriction phenotype is in agreement with a previous work reporting episodes of adaptive evolution (positive selection) in genes related with hypoxia tolerance in Cetacea, including genes involved in oxygen transport and regulation of vasoconstriction [35]. In addition, by expanding their analysis to other non-aquatic hypoxic environments, such as underground tunnels, they uncovered convergent evolution scenarios in species adapted to diving and burrowing [35]. A similar convergence is observed in the present work. In fact, in the mole *C. asiatica*, DRD_5_ was also predicted non-functional. Although this was the single DRD_5_ loss example found outside Cetacea, one cannot discard the putative contribution of alternative molecular events (i.e. post-translational mechanisms) towards trait loss [24], or event distinct physiological adaptations to overcome oxygen deprivation.

Besides diving physiology, DRD_5_-dependent sympathetic tone alterations could also contribute to the idiosyncratic sleep behavior observed in Cetacea [30,38]. Several physiological adjustments occur during sleep, encompassing thermoregulation, as well as endocrine, immune, pulmonary and cardiovascular functions [39]. In most mammals, sleep states lead to a decrease of the sympathetic tone, inducing vasodilation and decreasing blood pressure [39]. Thus, DRD_5_ loss could prevent sympathetic tone decrease in resting states paralleling the unihemispheric sleeping behavior and long-term vigilance observed in Cetacea.

Overall, our findings provide evidence for natural occurring KO for DRD_5_. Besides highlighting a molecular signature for vasoconstriction and blood pressure regulation in Cetacea, naturally occurring DRD_5_ KO could also provide useful frameworks to gain insight into hypertension and heart failure-induced peripheral vasoconstriction responses in humans [40,41].

## 4. Materials and Methods

### 4.1. Synteny analysis

To build the synteny maps for the DRD_5_ gene locus in Cetacea and *H. amphibius* (Figure 1) several annotated Cetacea genome assemblies were inspected and scrutinised using the NCBI browser, namely *O. orca* (GCF_000331955.2), *L. obliquidens* (GCF_003676395.1), *T. truncatus* (GCF_001922835.1), *D. leucas* (GCF_002288925.1), *N. a. asiaeorientalis* (GCF_003031525.1), *L. vexillifer* (GCF_000442215.1), P. *catodon* (GCF_002837175.1) and *B. a. scammoni* (GCF_000493695.1). *B. taurus* (GCF_002263795.1), a fully terrestrial relative of extant cetaceans, was used as reference. Next, (1) if DRD_5_ was annotated, the following procedure was used: five flanking genes, considering only the ones characterised and tagged as protein coding, from each side of DRD_5_ gene, were collected. (2) If DRD_5_ gene annotations were not found, *B. taurus* DRD_5_ direct flaking genes were used as reference genes to collect the corresponding flanking genes. In detail, CPEB2 was used as an anchor to search the rightwards DRD_5_ flanking genes, and OTOP1 to search the leftwards genes. Regarding *H. amphibius*, the synteny map was built via blast searches against the assembled, fragmented and not annotated genome of the same species, available at NCBI (GCA_002995585.1). *B. taurus* DRD_5_ flanking genes were used as reference, and using the discontiguous megablast task from blastn, the best blast hit (the hit with the highest alignment identity and query coverage) was retrieved and the coordinates of alignment in the target genome carefully inspected. *H. amphibius* synteny map was then built by sorting the genes according to the subject alignment coordinates within genes aligning at the same genomic scaffold.

### 4.2. Sequence retrieval and gene annotation

To perform manual gene annotation, all genomic regions of DRD_5_ sequences tagged as LQ were directly collected from NCBI. Regarding cetacean species with annotated genomes but without DRD_5_ gene annotations *(L. vellixifer and B. a. scammoni), the genomic sequence ranging from the upstream to the downstream flanking genes was collected.* For non-cetacean mammals, with annotated genomes but without DRD_5_ annotations two procedures were followed. (1) For each species, two DRD_5_ direct flanking genes were selected from a reference phylogenetic close species displaying non ‘low-quality protein’ DRD_5_ annotation (see Supplementary Table 2) and the test species region flanked by the same reference genes was further collected. (2) If the same flanking region exhibited severe fragmentation (Ns), using the same phylogenetic close species (Supplementary Table 2) DRD_5_ coding sequence as query, blastn searches were conducted against the Whole Shotgun Contigs dataset of the corresponding species via discontiguous megablast task. The genomic sequence corresponding to the blast hit with the highest alignment identity and query coverage amongst the others was retrieved.

For cetacean species with no annotated genomes *(B. bonaerensis, E. robustus, B. mysticetus, S. chinensis), as well as H. amphibius, genomic sequences were retrieved through blastn searches in the corresponding genome assembly using the B. taurus DRD5 coding sequence as query. The best genomic scaffold considered for each species corresponded to the blast hit with the highest query coverage and identity value amongst total hits. Due to the presence of a fragmented genomic region concerning T. truncatus DRD5 gene annotation, the same blast search procedure was executed for this species, using as target the Whole Genome Shotgun contig dataset of the same species. The gene annotation concerning this species was performed in the retrieved contig corresponding to the blast hit with the highest query coverage and alignment identity value.*

Collected genomic sequences were further imported into Geneious Prime 2019 (www.geneious.com) and the DRD_5_ gene coding sequences were manually annotated for each species. Briefly, using the built-in map to reference tool with the highest sensibility parameter selected, the reference single-exon DRD_5_ gene, 3’ and 5’ UTR flanked, was mapped against the corresponding genomic sequence of the in-study species and aligned regions carefully screened for ORF abolishing mutations including frameshift mutations and premature stop codons. For Cetacea and *H. amphibius DRD5 gene annotation, B. taurus DRD5 was selected as reference. Regarding non-cetacean mammals DRD5 annotation*, different references were chosen according to the phylogenetic relationships between reference and test species (Supplementary Table 2). The identified mutations were next validated using the sequencing run raw data (sequencing reads) retrieved from at least two independents genomic NCBI SRA projects (when available).

## Author Contributions

conceptualization, R.R. and L.F.C.C.; methodology, L.Q.A. and L.F.C.C.; investigation, L.Q.A., R.R. and J.A.; data curation, L.Q.A.; writing—original draft preparation, R.R., L.Q.A. and L.F:C.C.; writing—review and editing, All; supervision, R.R.; project administration, L.F.C.C.; funding acquisition, L.F.C.C.

## Acknowledgments

We acknowledge the various genome consortiums for sequencing and assembling the genomes.

